# WFD ecological status indicator shows poor correlation with flow parameters in a large Alpine catchment

**DOI:** 10.1101/419804

**Authors:** Stefano Larsen, Maria Cristina Bruno, Guido Zolezzi

## Abstract

Since the implementation of the Water Framework Directive, the ecological status of European running waters has been evaluated using a set of harmonised ecological indicators that should guide conservation and restoration actions. Among these, the restoration of the natural flow regime (ecological flows) is considered indispensable for the achievement of the good ecological status, and yet the sensitivity of the current biological indicators to hydrologic parameters remains understudied. The Italian Star_ICMi well represents other similar WFD indicators; it is a macroinvertebrate-based multimetric index officially adopted to assess the ecological status of running waters at the national level. Recent legislation has also included the Star_ICMi as one of the indicators used to assess and prescribe ecological flows in river reaches regulated by water abstraction. However, the relationship between river hydrology and the Star_ICMi index is so far virtually unknown. Using data from the Trentino - Alto Adige Alpine region, we first assessed the relationship between the Star_ICMi and synthetic descriptors of the physico-chemical (LIMeco) and morphological (MQI) status of respectively 280 and 184 river reaches. Then, we examined the relation between the Star_ICMi and a set of ecologically-relevant hydrologic parameters derived from discharge time-series measured at 21 hydrometric stations, representing both natural and regulated river reaches. Although the Star_ICMi showed significant and linear relationships with the physico-chemical character and, slightly, with the morphological quality of the reaches, its response to flow parameters appeared weak or non-existent when examined with linear models. Mixed quantile regressions allowed the identification of flow parameters that represented limiting factors for macroinvertebrate communities and the associated Star_ICMi scores. In particular, the index showed ‘negative floors’ where lower values were observed in reaches with large temporal variation in flow magnitude as well as frequent low and high flow events. The modelled quantiles also tracked the transition of the index from acceptable to unacceptable conditions.

The results suggest that while the central tendency of the Star_ICMi index is not strongly influenced by river flow character, some key flow parameters represent limiting factors that allow the index to reach its lowest values, eventually ‘pushing’ the site towards unacceptable ecological conditions. The identification of limiting flow parameters can aid the setting of hydrologic thresholds over which ecological impairment is likely to occur. Overall, however, results imply caution is needed in using biological indicator like the Star_ICMi for the quantitative assessment and design of ecological flows.

## 1. Introduction

The Water Framework Directive (WFD; European Commission, 2000) is the principal legislative framework concerning the management and protection of European waters. Through the definition of common approaches, the WFD requires Member States to achieve ‘good ecological status’ objectives for water bodies. Among the quality elements guiding the status classification of streams and rivers, the hydrologic regime (the quantity and dynamics of river flow, sensu Poff et al., 1997) is considered central in supporting the biological elements and thus the achievement of good ecological status. Although not explicitly mentioned in the WFD, the concept of ‘ecological flow’ (E-flow) is increasingly considered and implemented in many river basin management plans. Within the EU context, E-flows are defined as “an hydrologic regime consistent with the achievement of the environmental objectives of the WFD in natural surface water bodies”, and specific recommendations on the definition of E-flows and their use in status assessment were also recently provided (Guidance 31 by the European Commission; WFD CIS, 2015). In particular, the Guidance 31 states that the “Ecological impacts of hydrological alterations and their significance should be ultimately assessed with biological indicators built on monitoring data that are specifically sensitive to hydrological alterations”.

The use of biological indicators has a long tradition in freshwater ecology where fish and macroinvertebrate based indices are widely used to define the ecological integrity of waterbodies (e.g. De Pauw et al., 2006; Rosenberg and Resh, 1993). In Europe, after the implementation of the WFD, there has been substantial effort to harmonise the different eco-bio-indicators across EU Countries (e.g for macroinvertebrates: Hering et al., 2004; Verdonschot and Moog, 2006). These indicators are used to define the ecological status of running waters and guide conservation and restoration effort.

However, although river organisms are clearly influenced by the hydrology, most present bioindicators were developed to emphasise organisms sensitivity to organic pollution and habitat degradation, and hence appear rather insensitive to hydrological alterations (Friberg, 2014; Poff and Zimmerman, 2010). Although some countries developed hydrologically-sensitive indicators based on flow preference of benthic invertebrates (UK: Extence et al., 1999; NZ: Greenwood et al., 2016), these are not yet implemented in the WFD. The implementation of evidence-based E-flows should be based on a sound understanding of the relation between river ecology (e.g. biodiversity) and flow characteristics (flow-ecology relationship: Rosenfeld, 2017; Stewart-Koster et al., 2014), ultimately requiring a fundamental association between water quantity and ecological quality. Yet, more effort has been dedicated internationally towards the definition and modelling of E-flows and water allocation for regulated rivers (e.g. residual flow) compared to the quantification of flow-ecology relationships (Davies et al., 2014; Tonkin et al., 2014). The natural flow paradigm is at the heart of the E-flow concept in that modified flow regimes should incorporate the natural variability in terms of flow magnitude, frequency, duration, timing and rate of change (Poff et al., 1997). Since the publication of the Nature Conservancy’s Indicator of Hydrologic Alteration (IHA; Richter et al., 1997), parameters quantifying the different components of the flow regime have been widely used to characterise natural flow regimes and its alterations as well as to identify ‘ecologically relevant’ hydrological drivers (e.g. Worrall et al., 2014). Similarly, flow-ecology studies quantifying the influence of individual flow parameters on in-stream communities have been flourishing steadily in recent times (Tonkin et al., 2014); however, those that specifically assessed the response of multi-metric indicators such as those adopted by the WFD are scarce (Belmar et al., 2018; Monk et al., 2006; Nebra et al., 2014). This is surprising considering the emphasis given by the WFD on water abstraction, ranked as the second most common pressure on EU water bodies (WFD CIS, 2015). Therefore, assessing how current WFD biological indicators respond to the different components of the flow regime is a prerequisite for managing E-flows and developing more specific indicators.

Here we used a framework based on flow-ecology relationship to investigate the performance of a WFD bio-indicator to characterise hydrologic regimes and their alterations. As a case study, we used the macroinvertebrate-based Star_ICMi (Buffagni and Erba, 2007) officially adopted by the Italian legislation as the Biological Quality Element to guide the classification of running waters according to the WFD. The index is based on six normalized and weighted metrics also adopted by other EU Countries (Buffagni et al., 2006), and includes taxonomic richness and diversity, as well as taxa sensitivity to organic pollution. Alongside other purely hydrological and habitat-based eco-hydraulic indicators, the Star_ICMi also represents one of the methods adopted by the Italian law for the determination of E-flows in regulated rivers. However, since its official introduction in Italy, the few available studies have indicated a rather low sensitivity of the Star_ICMi to discharge alterations, especially where these are not coupled with a deterioration of water-quality, as it often occurs in Alpine and perialpine streams affected by hydropower regulation (Laini et al., 2018; Quadroni et al., 2017; Salmaso et al., 2018). Recently, some critical issues in using the STAR_ICMi to determine E-flows have been raised based mainly on the apparent lack of a direct relationship between the index and river discharge (Spitale and Bruno, 2018). Surprisingly, despite the specific requirements of the WFD, so far the relationship between the Star_ICMi and different flow parameters describing river discharge has been virtually unexplored in Italy (but see Laini et al., 2018). However, investigating how this ecological quality indicator responds to flow characteristics is indispensible to evaluate its use within the context of E-flows.

As a representative case study for the Italian Alpine area we analysed data from the Trentino-Alto Adige region where the main alterations of the natural flow regime are essentially related to hydropower schemes (Zolezzi et al., 2009). The study has two main objectives: first, we used the region-wide dataset to investigate the responses of the STAR_ICMi to the physico-chemical and morphological character of river reaches, as described by synthetic WFD quality elements. Second, by identifying a set of monitoring stations for which river discharge time-series were available, we quantified the relationship between the Star_ICMi and a set of ecologically-relevant flow parameters, using both linear and quantile regressions. Because the latter analysis was based on limited data points, we did not attempt to disentangle and rank the individual effect of multiple environmental factors besides hydrologic regime (e.g. as in Booker et al., 2015). Instead, we appraised the extent to which other environmental covariates (i.e., anthropogenic stressors) might have influenced the observed flow-ecology relationship using the Procrustes analysis. Specifically, we tested if the correlation between hydrological parameters and macroinvertebrate communities increased with altitude where the influence of other anthropogenic stressors (e.g. nutrients, local land use) appeared to be weaker.

## 2. Methods

### 2.1 Study area

The Trentino-Alto Adige is a region in Northeast Italy with a surface of c. 13.000 Km^2^ and a population of c. 900.000 inhabitants. The region mostly lays within the Alps with more than 75% of its territory above 1000 m of altitude. The Adige River and its tributaries form the largest river basin occupying about 80% of its territory. Minor river basins in the region included in the study were the Sarca, Brenta, Chiese and Vanoi. A total of 280 study reaches were included, which form the monitoring network of the Environmental Protection Agencies of the Provinces of Trento and Bolzano, across an altitudinal range of 175 - 1800 m a.s.l (Fig. 1).

**Figure 1.**
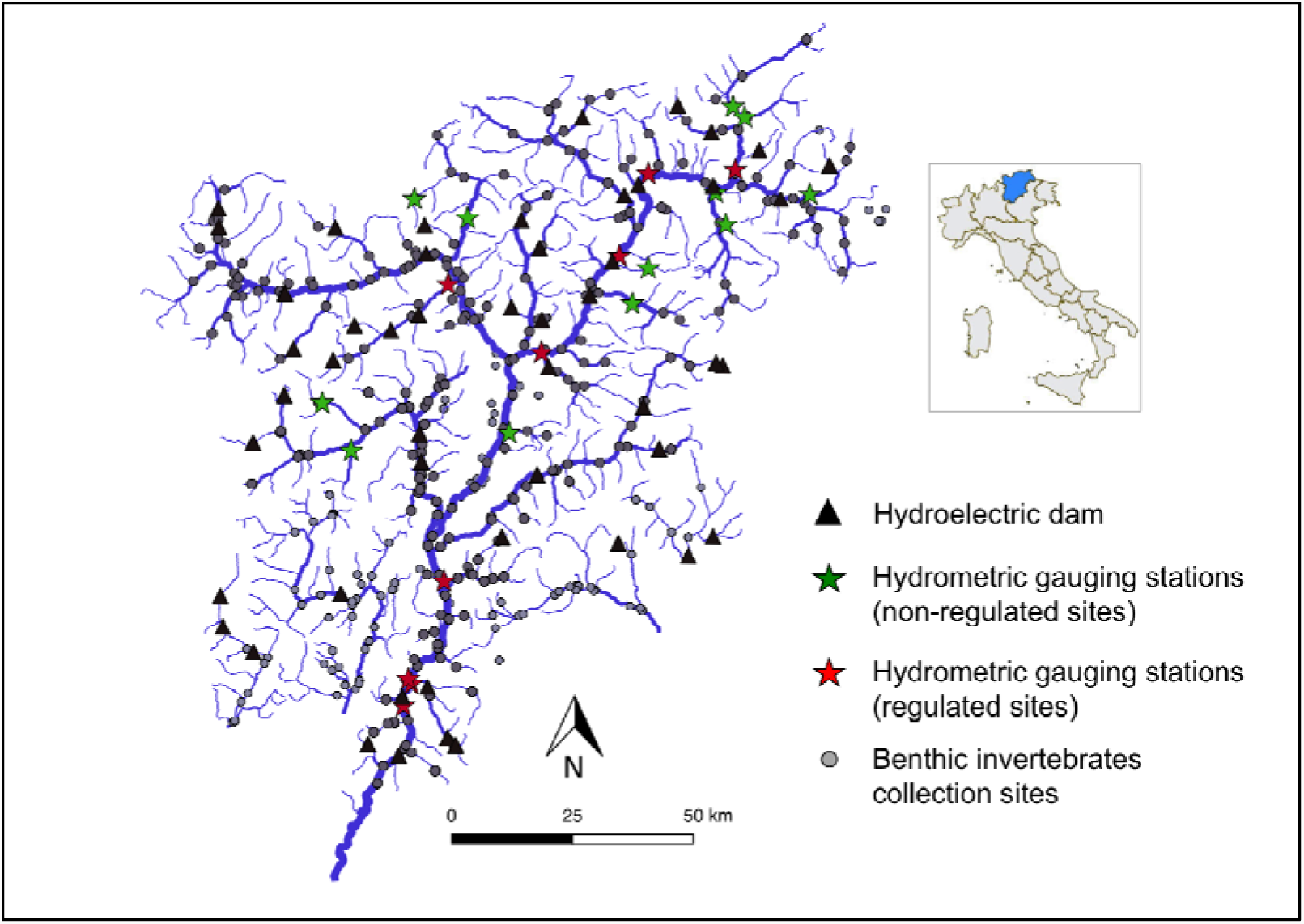
Map of the main river networks in the Trentino-Alto Adige region in NE Italy and the analysed gauging stations (stars) and biological sampling stations (dots).

### 2.2 Data collection and computation of the WFD indicators

Data used in the present study come from the institutional monitoring programme of the Environmental Protection Agencies of the Provinces of Trento and Bolzano. Benthic macroinvertebrates were collected in 280 stream reaches between 2009 and 2014 (Fig. 1). Sampling followed the multi-habitat proportional technique according to the AQEM protocol (Hering et al., 2004), in which 10 replicate Suber samples (0.1m^2^) were distributed along the reach proportionally to the different micro-habitat types present. Specimens were identified to genus and family levels as required for the calculation of the Star_ICMi. The index is computed combining sub-metrics related to the tolerance, richness and diversity of the different macroinvertebrate taxa observed (Appendix A in Supplementary Material).

In most of these biological sampling sites, data for the formulation of two additional WFD indicators were also gathered. To assess the physico-chemical quality element, we used the LIMeco index (“Livello di Inquinamento dai Macrodescrittori per lo stato ecologico”), which scores river water quality in terms of dissolved oxygen and nutrient concentration (Azzellino et al., 2015), with data for 280 reaches. The morphological quality was assessed with the Morphological Quality Index (MQI; Rinaldi et al., 2013), with data available for a subset of 184 reaches. The MQI provides a score to the morphological quality of a river reach based on three main elements: geomorphological functionality (accounting for longitudinal and lateral continuity of river processes, channel patterns, river bed structure and substratum, riparian vegetation), degree of artificiality (e.g. presence of local and remote sources of hydro-morphological alterations, such as sediment mining, levees and embankments, artificial reservoirs), and observed recent channel adjustments.

Hydrological information was available from gauging stations located along the Adige River network (managed by the Ufficio Dighe for the Autonomous Province of Trento, and by the Ufficio Idrografico for the Autonomous Province of Bolzano), and we selected 21 gauged stations (Table 1) in proximity to biological sampling site (<5km stream distance, no influence of major tributaries) so as to pair hydrological and ecological data (Fig. 1). Overall discharge time-series differed in length among stations, but continuous flow records were available for all stations from 2007. This allowed us to associate 1-year antecedent flow series with each biological sample.

**Table 1.**
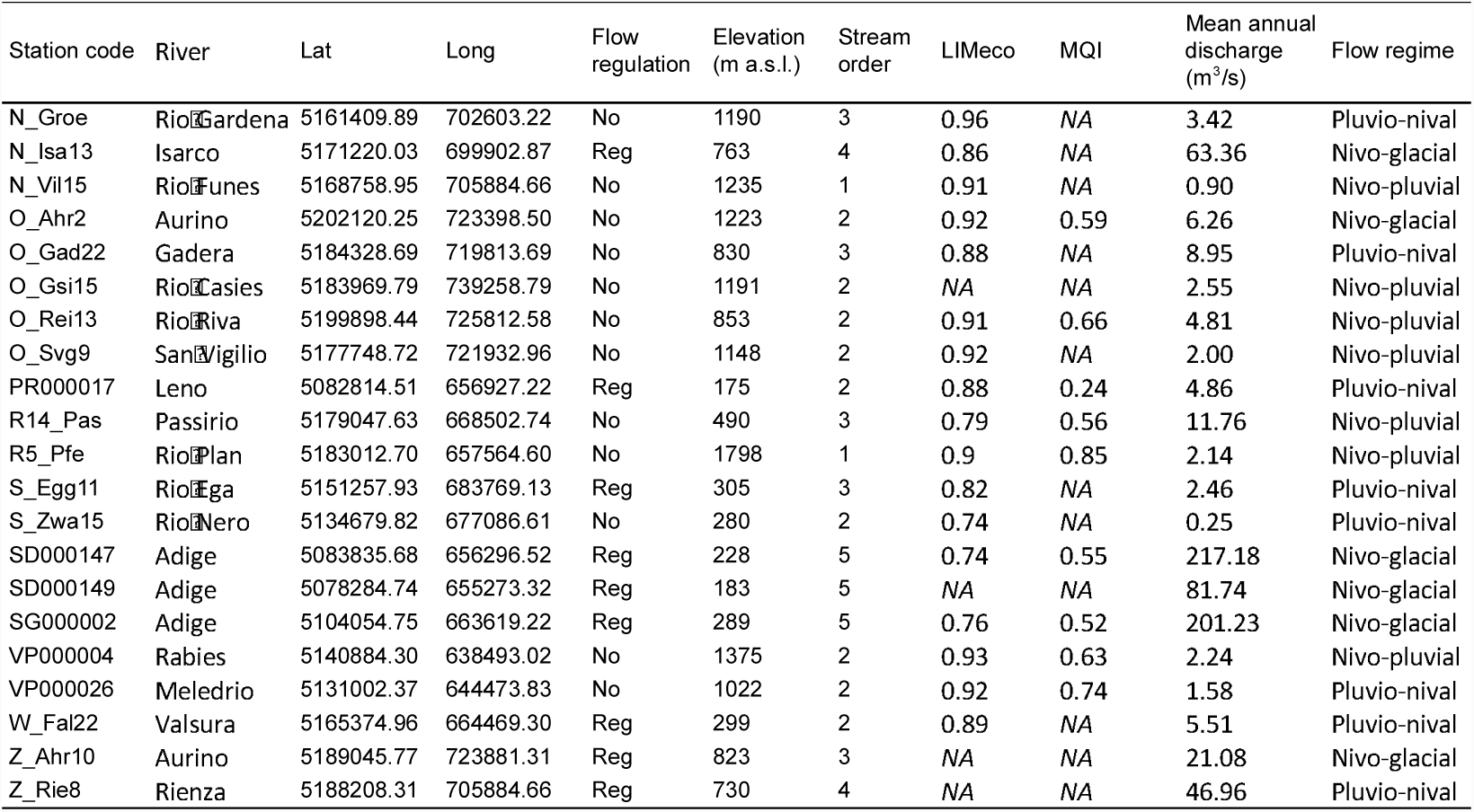
Main characteristics of the hydrometric gauging stations used to derive flow parameters from discharge time-series.

### 2.3 Data analyses

Across the 280 monitoring sites, macroinvertebrate sampling occurred multiple times between 2009 and 2014 (3-10 times per site). We therefore calculated the mean Star_ICMi value to characterise the biological quality of each site. Similarly, multiple values of the LIMeco and MQI indices were averaged per site. For the first goal (relation among WFD quality indicators), we used ordinary least square regressions to relate the Star_ICMi index with the LIMeco and MQI indices.

For our second goal (flow-ecology relationship), we used the available discharge time-series to derive 21 flow parameters (Table 2) based on daily flow values (normalised relative to annual means). Following previous studies (Belmar et al., 2018; Worrall et al., 2014) two temporal scales were considered: 1-year and 60-days preceding the collection of benthic macroinvertebrates, thus representing the influence of both long and short-term antecedent hydrologic conditions. Flow parameters were derived following Indicator of Hydrologic Alteration approach (IHA; Richter et al., 1997) using the ‘IHA’ implementation in R (R Core Team, 2017). The flow parameters represented ecologically relevant hydrologic characteristics regarding magnitude (e.g. 1-7-90 days maximum and minimum flow), frequency and duration (e.g. number and duration of low and high pulses), rate of change and variation (e.g. rise and fall rates, CV). No automatic selection of flow parameters or synthesis was performed (e.g. PCA reduction), so as to avoid the exclusion of relevant parameters, and to facilitate the interpretation of results (e.g. Schneider and Petrin, 2017). In addition, the overall number of parameters included was small (compared to most publication where >100 metrics are used) and represented arguably the minimum set of ecologically relevant flow characteristics. Parameters related to the timing of flow events were not calculated, because the computation would require longer (multiple-years) flow time-series, and because macroinvertebrate collection was conducted over different months of the year. Star_ICMi values from repeated observations in time (multiple biological samples per site) were not averaged in this case, but were all included for a total of 80 samples, each paired with 1-year hydrological information. This allowed us to increase statistical power and aid the visual interpretation of complex relationships. The longitudinal structure of the data was accounted for by including ‘site’ as random factor in linear mixed-models relating the Star_ICMi to flow parameters, using the nlme package in R (Pinheiro et al., 2018). The proportion of variance explained by the fixed factors (i.e. flow parameters) was expressed as marginal R^2^ using the r.squaredGLMM function in the MuMin package (Bartoń, 2018).

**Table 2.**
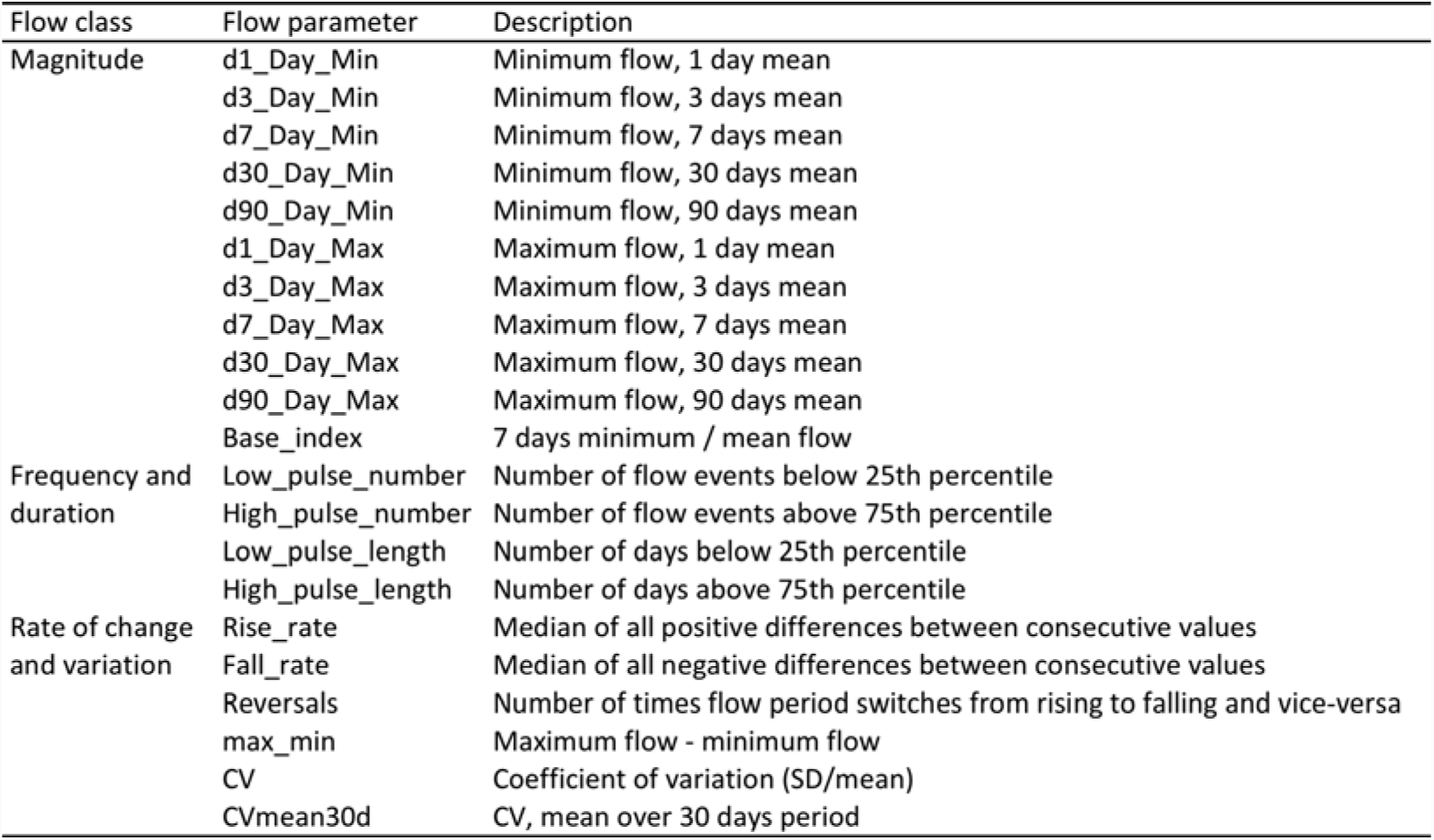
Flow parameters derived from daily flow-series included in the analyses

The 21 gauged reaches represented rivers with natural flow regime as well as reaches regulated by upstream hydropower schemes. These were all included in the analyses, because i) we were interested in assessing the relation between the Star_ICMi and specific flow parameters rather than quantifying differences among river reaches and ii) we wanted to include the full range of flow parameters expected in the region. Nonetheless, for aiding visual interpretation, regulated and non-regulated reaches were differently identified in the plots.

In our analytical approach, we recognise that streamflow conditions are among the many factors that influence macroinvertebrate assemblages across the study sites. These include, for instance, water quality parameters and temperature, resource availability and riverbed morphology among others (Allan, 1995). However, streamflow characteristics in some reaches can represent a limiting factor for macroinvertebrates, where other stream features would allow different density or diversity to be observed. These limits can be considered as either ‘ceilings’ or ‘floors’ when the biological metric shows upper or lower limits as a function of a flow parameter, respectively. In these cases, the biological metric is unlikely to display a central response to flow parameters and ordinary regressions are not suited to quantify the limits (Konrad et al., 2008; Lancaster and Belyea, 2006). Conversely, quantile regressions allow modelling the effect of a predictor variable over different quantiles of the dependent variable. In other words, the model fits the ‘limiting response’ of the *y* variable by identifying its conditional quantiles with respect to the predictor variable *x*. When quantifying the 80th percentile, for example, 80% of the values of *y* are equal or less than the modelled function of *x* (Cade and Noon, 2003). In the present study, we assessed both the central and limiting response of the Star_ICMi index to the different flow parameters. We used linear mixed-models to account for repeated sampling within site using the ‘nlme’ package in R. For the quantile approach we employed a recently developed algorithm for linear quantile mixed-models implemented in the ‘lqmm’ package in R (Geraci, 2014).

Lastly, we used Procrustes analysis to appraise the extent to which other confounding factors may influence the flow-ecology relationship in the study area. Similarly to Mantel test, Procrustes analysis quantifies the association between multivariate data matrices, but it also provides a vector of residuals that represent the differences between homologous observations (i.e. sites, samples) across the two matrices (Lisboa et al., 2014). The residuals vector is a measure of the fit between the two matrices and can be used to further understand how the matrices are related. For example, the Procrustes residuals can be used to appraise whether another factor influenced the degree of matching between observations. Here we used Procrustes analysis to quantify the match between the matrix of macroinvertebrate communities (*samples × taxa*) and the matrix of flow parameters (*samples × parameters*). Then, we extracted the residuals vector and used it to investigate the influence of other environmental covariates. Specifically we used altitude (ranging 175 - 1800 m a.s.l.) as a proxy for many correlated factors and stressors such as temperature, land-use and water quality, and a linear mixed-models was used to relate the Procrustes residuals vector with altitude. Procrustes analysis requires the same dimensionality between multivariate matrices. Therefore, we first harmonised the dimensionality of each matrix using Principal Component Analysis (PCA) by keeping the first five components for each matrix. Macroinvertebrate densities were Hellinger-transformed prior to PCA (Lisboa et al., 2014).

## 3 Results

The Star_ICMi index showed linear and relatively strong (R^2^=0.36, P <0.0001; n=280) correlations with the LIMeco index and, to a lesser extent, with the MQI (R^2^=0.2, P<0.0001; n=184) (Fig. 2).

**Figure 2.**
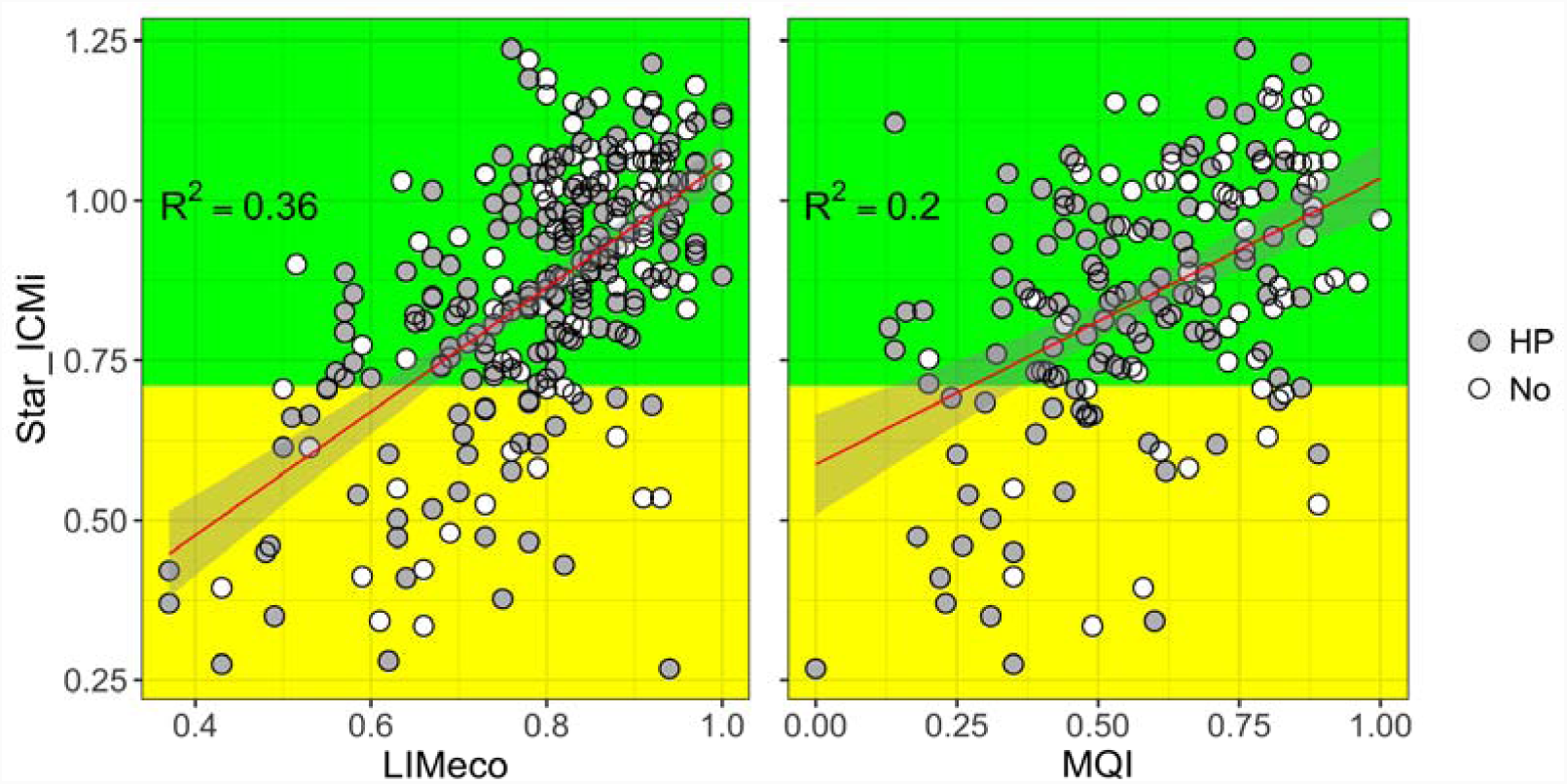
Regression of the Star_ICMi with LIMeco and MQI. Red lines = linear fit; grey areas = 95% CI. Background colours denote threshold between acceptable (green) and unacceptable (yellow) conditions (sensu WFD). Grey circles = reaches affected by hydropower upstream (HP); white circles = reaches not affected by hydropower (No)

Only three of the flow parameters calculated from 1-year flow series were significantly (at P<0.05) and negatively correlated with the Star_ICMi according to linear mixed models, namely CV_mean30d (marginal R^2^=0.3), d1_Day_Max (marginal R^2^=0.1) and max-min (marginal R^2^=0.1) (Fig. S1 in SM; see description of the parameters in Table 1). No significant correlations were observed when flow parameters were derived from 60-days flow series preceding the macroinvertebrates collection (Fig.S2 in SM).

Conversely, the use of quantile regressions allowed the identification of additional flow parameters that appeared to limit the scores of the Star_ICMi (Fig. 3). In particular, peak flows (d1_Day_Max), the number of low and high pulses as well as parameters related to flow variation (CV, max_min) represented ‘negative floors’, which led the Star_ICMi below unacceptable conditions. Put in other words, lower values of the aforementioned flow parameters limited the minimum scores of the Star_ICMi.

**Figure 3.**
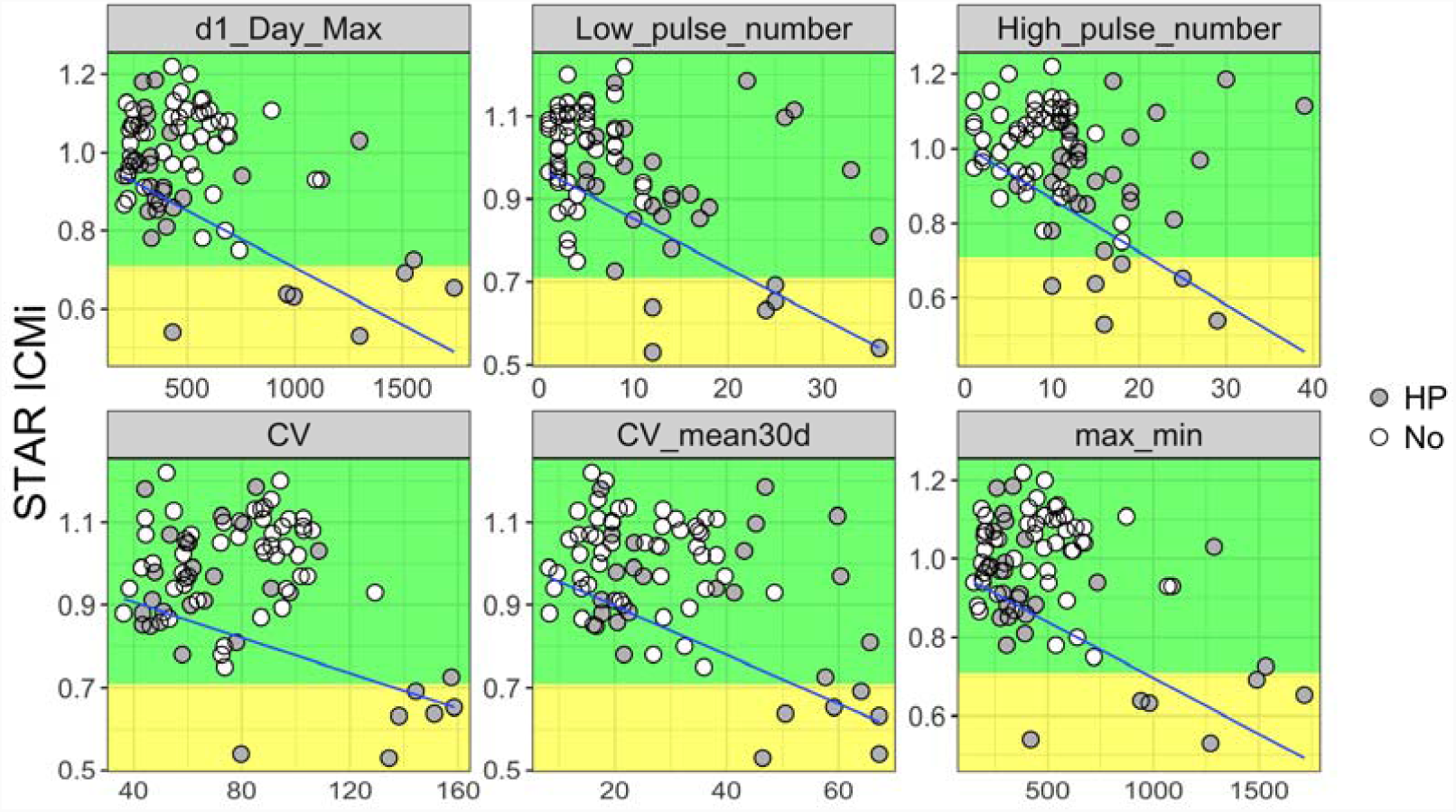
The Star_ICMi vs IHA metrics based on flow time-series from 1-year preceding the biological sampling. Blue line indicates significant mixed quantile regression at q=0.2. Significance levels are at P<0.01 for all parameters except for CV at P=0.04. Background colours denote threshold between acceptable (green) and unacceptable (yellow) conditions (sensu WFD). Grey circles = reaches affected by hydropower upstream (HP); white circles = reaches not affected by hydropower (No)

When flow parameters were derived from 60-days flow time-series, the influence of low and high flow pulse number remained significant, while the effect of hydrologic reversals became apparent, also in the form of a ‘negative floor’ (Fig. 4). That is, the minimum scores of the Star_ICMi were observed in reaches characterised by frequent hydrologic reversals. At this shorter time-scale, the effect of daily rate of change in flow also became apparent, with a ‘negative floor’ observed with Rise_rate (Fig.4).

**Figure 4.**
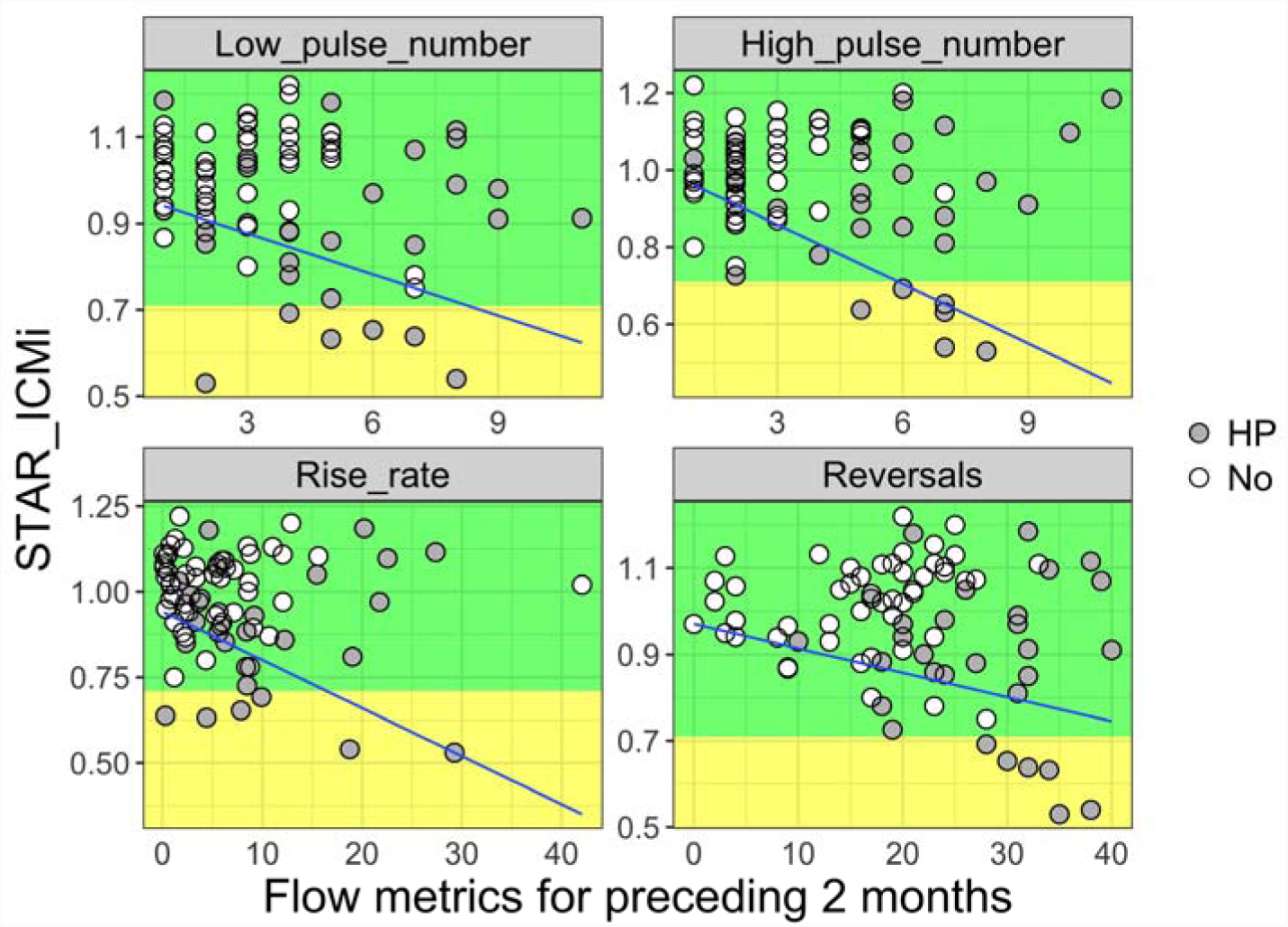
Star_ICMi vs IHA metrics based on flow series from 2 months preceding the biological sampling. Blue line indicates significant quantile regression at q=0.2. Significance levels are at P<0.05. Background color denotes threshold between acceptable (green) and unacceptable (yellow) conditions (sensu WFD). Grey circles = reaches affected by hydropower upstream (HP); white circles = reaches not affected by hydropower (No)

Our study reaches encompassed a wide altitudinal range, which potentially acted as a confounding factor in the analysis of flow-ecology relationships as altitude was related to changes in the main physico-chemical parameters and local land use across the study reaches (Fig. S3 in SM). In fact, the Star_ICMi index increased linearly with altitude (Fig 5A; mixed-model marginal R^2^=0.2, P=0.01), and most of the regulated river reaches occurred at lower elevations. The residuals vector from the Procrustes analysis associating the matrices of flow parameters and macroinvertebrate communities declined with altitude (Fig. 5B; mixed-model marginal R^2^=0.15, P=0.04), indicating a significant and negative effect of altitude. These results indicate that the match between the biota (taxonomic matrix) and the hydrological parameters was stronger in higher-altitude locations, likely because the influence of confounding stressors was weaker.

**Fig. 5.**
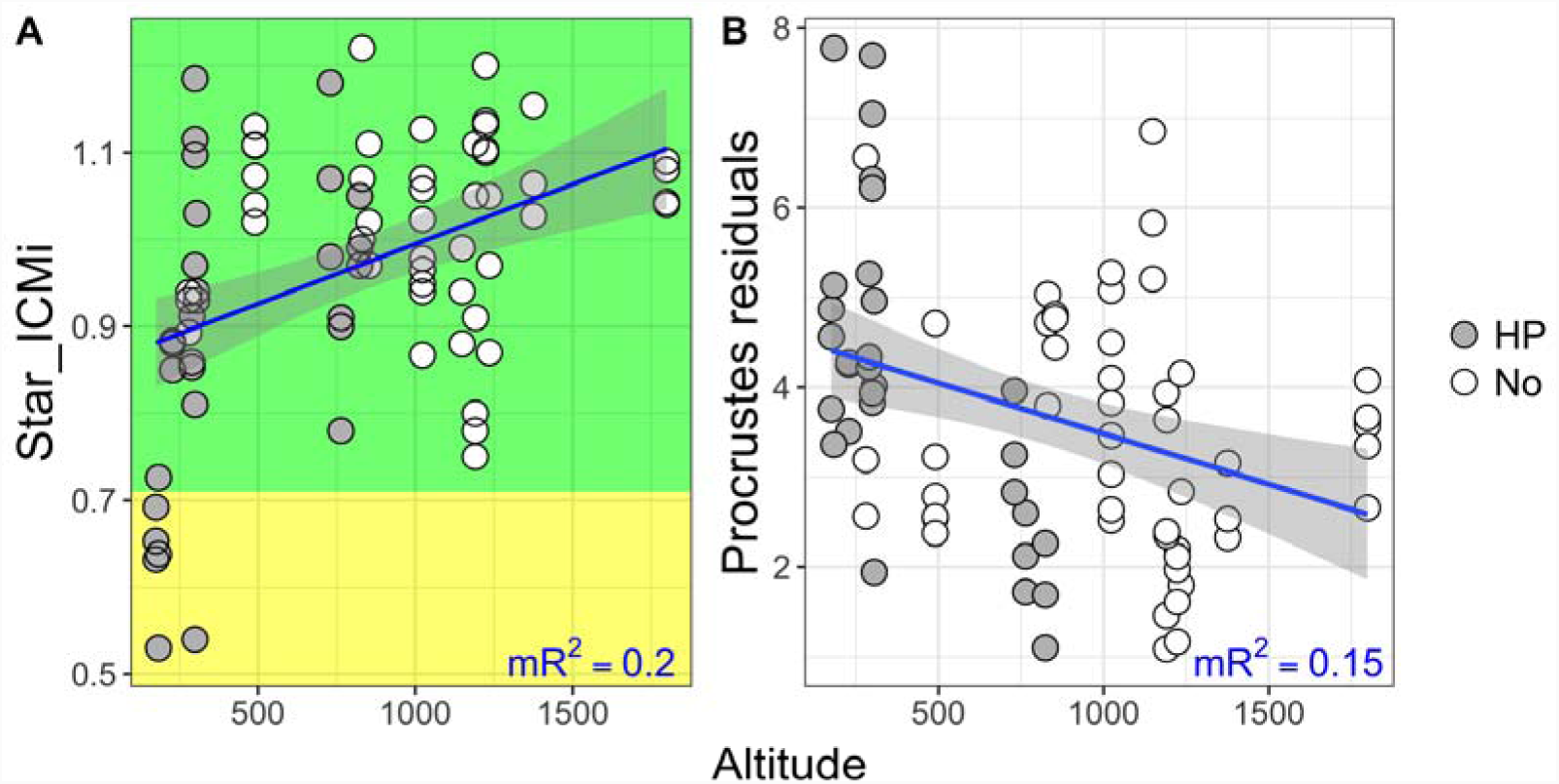
Relation between the Star_ICMI (A) and Procrustes residual (B) with altitude. Legend as in Fig. 3 and 4.

## 4. Discussion

We used monitoring data from a large Alpine river network to assess the relationship between WFD quality elements, and the response of the official Italian macroinvertebrate-based indicator to measured hydrological parameters.

The Star_ICMi showed a relatively strong relation with the physico-chemical character of the river reaches, as expressed by the LIMeco index. This was expected, as stream macroinvertebrates are known to be particularly sensitive to water quality parameters (e.g. Friberg et al., 2010; Guilpart et al., 2012). Moreover, the LIMeco index specifically reflects the concentration of organic pollutants (nitrates and phosphates), to which the Star_ICMi is designed to respond (Quadroni et al., 2017). Parallel findings were recently reported by Azzellino et al. (2015) for the nearby Lombardy region (northern Italy), where water quality, as expressed by the LIMeco scores, explained *c.* 50% of the variation in the Star_ICMi across a range of river reaches with similar environmental settings as those studied here.

The influence of rivers’ morphological features on benthic invertebrates, as expressed by the MQI, was also significant, but apparently weaker. The MQI reflects the integrity of both channel and riparian habitat that can influence in-stream organisms in multiple ways, for instance by providing refugia and resources (Matthaei et al., 2000; Naiman and Décamps, 1997). Hence, our results suggest that across the Trentino-Alto Adige region, physico-chemical water quality was the main determinant of macroinvertebrate community integrity as measured by the STAR_ICMi, which was only secondarily affected by riparian and in-stream morphological features. Other studies in Italy and elsewhere indicated that substratum and riparian characteristics mostly influenced the functional composition (e.g. feeding habits) of benthic invertebrates, rather than their taxonomic identity and diversity (e.g. Larsen and Ormerod, 2010; Manfrin et al., 2016). This could in part explain the lower sensitivity of the taxonomic-based Star_ICMi to the MQI.

Results from our second objective also indicated a rather poor sensitivity of the Star_ICMi index to hydrological parameters. Analyses of flow-ecology relationship were based on a reduced sample size, because we selected those reaches for which daily discharge time-series were available from adjacent gauging stations. Nonetheless, the reaches were distributed across the whole extent of the study area and included both natural and regulated rivers and likely represented the entire range of flow parameters observed in the region. These parameters were derived at two temporal scales (1-year and 60-days preceding biological sampling) and provided similar but not identical results. When assessing the central response (using linear mixed-models), only parameters derived at 1-year time scale appeared to significantly and negatively affect the biological indicator, and included the monthly coefficient of variation in flow and the overall range and maximum daily flow. These all indicate a generally negative effect of large daily flow variation on aquatic communities, as also reported elsewhere (e.g. Bruno et al., 2010; Konrad et al., 2008; McGarvey, 2014).

It is important to highlight how focusing on the central response of macroinvertebrate metrics can provide only limited insight into the effects of streamflow. Lotic invertebrates are influenced by a wide range of abiotic and biotic factors and are unlikely to display a central or linear response to streamflow parameters (Konrad et al., 2008; Rosenfeld, 2017). In these cases, the use of quantile regressions allow the identification of those factors that appear to locally limit the maximum or minimum values of the response variable. Previous studies with benthic invertebrates have shown the validity of this approach for the identification of the environmental constraints on local community density and richness (Fornaroli et al., 2015; Lancaster and Belyea, 2006). Here, we used a novel approach based on mixed quantile models and were able to identify some key flow parameters that appeared to determine the lower limits of the Star_ICMi. Interestingly, the significant quantiles all took the form of ‘negative floors’, whereas no significant ‘ceilings’ were observed. Negative floors imply that the lower limits of the biological indicator declined with increasing values of the flow parameters. This resulted in the modelled quantiles to apparently ‘track’ the transition of the ecosystem into unacceptable conditions. Viewed in terms of ecological constraints, these negative floors suggest that the biological integrity was maintained within acceptable conditions (sensu WFD) by lower values of the flow parameters, which evidently represented favourable hydrologic conditions. As the value of the flow parameters increased, the hydrologic conditions deteriorated thus leading some sites to drop to a lower quality status. The identified limiting parameters were mostly related to flow variation and the frequency of flow events. Specifically, we found that high peak flows, frequent low and high pulses and large variations in daily flow (as CV) apparently acted as stressors for macroinvertebrates leading to a marked decline in the Star_ICMi at some sites. Similar patterns were observed by Konrad et al. (2008) in 111 stream sites in the western U.S.A., where invertebrate abundance and the proportion of intolerant taxa showed quantile relations in the form of negative floors with parameters describing discharge variation.

Our results also parallel those Worrall et al. (2014) in showing that not only the long-term flow regime, but also short-term antecedent flow conditions can influence benthic communities. Our modelling procedure identified additional flow parameters as limiting factors when derived from 60-days preceding macroinvertebrate collection. In this case, the lower values of the Star_ICMi were limited by large daily rise rates in flow and the number of hydrologic reversals, which are also parameters quantifying hydrologic variations. It should be noted that most reaches characterised by higher values of the limiting flow parameters (e.g. yearly CV, frequent high and low pulses, reversals) were located downstream of hydropower plants, albeit at different distances. Hydropower operations in the region are known to affect the natural flow variability by often increasing the frequency and amplitude of flow oscillations and sharp transitions (Zolezzi et al., 2009). Therefore, the identified limiting flow parameters were likely outside their natural range of variability, and thus represented stressing factors for the communities.

The large scatter or variance in the relation between the Star_ICMi and flow parameters clearly indicates the influence of additional limiting factors (e.g. water quality and altitude, as seen here). Disentangling the different source of variation in these cases can be challenging as these can include both natural and anthropogenic factors as well as biotic and abiotic processes. In these cases, the use of quantile modelling has offered clear advantages (Fornaroli et al., 2015; Konrad et al., 2008), as also observed in the present study. We specifically attempted to quantify the influence of other covariates on the observed flow-ecology relationship using the multivariate Procrustes analysis and the associated residuals vector. We used Procrustes to first associate the matrices of flow parameters and macroinvertebrate communities (sites *x* taxa densities). Then, we derived the vector of residuals that quantified the mismatch between homologous observations (sites) in the multivariate space defined by the two matrices (Lisboa et al., 2014). This residuals vector showed a significant and negative correlation with altitude. This means not only that altitude acted as an important covariate, but also that the match between the biota and the hydrology was stronger in upland reaches compared to lowlands. Upland reaches were likely less influenced by potential confounding factors related to human activity, including nutrient inputs and land use conversion, and the influence of streamflow characteristics on local communities was evidently stronger.

This contingency has wider implications for the development of general flow-ecology relationships in the area and potentially across Europe, where analogous biological indicators are adopted in line with the WFD requirements (Buffagni et al., 2006), and further emphasises the need for research and management that acknowledges the complexity of multiple stressors acting on river ecosystems (Ormerod et al., 2010).

## 5. Conclusions

In this study, we used data from a large Alpine river network to assess the relation among different WFD quality elements and to contribute to the development of evidence-based ecological flows. This was also motivated by the warning from the European commission (WFD CIS, 2015) stating that “in cases where hydrological alterations are likely to prevent the achievement of environmental objectives, the assessment of the gap between the current flow regime and the ecological flow is a critical step to inform the design of the programme of measures”.

Our results suggest that existing macroinvertebrate-based biological indicators, like the Star_ICMi used as a case study and prescribed by the Italian national legislation, may mostly reflect local physico-chemical water quality and to a lesser extent the morphological integrity of the reaches, as expressed here by the LIMeco and MQI descriptors, respectively. This result was expected given previous observations and the known sensitivity of the index to organic pollution (Azzellino et al., 2015; Quadroni et al., 2017). As such, the Star_ICMi showed rather poor correlations with flow parameters when examined in its central response. Nonetheless, quantile modelling allowed the identification of key flow parameters that limited the minimum scores of the index (i.e. ‘negative floors’), and apparently tracked the transition of the ecosystem into unacceptable conditions. The identification of these flow limits can aid the implementation of E-flows by allowing managers to compare local conditions with the given limits and set hydrologic thresholds over which ecological impairment is likely to occur. In the study area, most of the negative limits identified were related to the magnitude and frequency of flow variations, which were likely altered by upstream hydropower operations. However, as also emphasised by Konrad et al. (2008), these limits show that the biological response to local hydrologic characteristics is contingent upon a range of local and regional factors including both natural (e.g. altitude) and anthropogenic ones (water quality), as demonstrated here by the Procrustes analysis. This has clear implications for both fundamental flow-ecology research and for water management, because the response of biological communities and associated indicators to flow regulation cannot be predicted without detailed information on the wider environmental setting of a river reach.

Although some important limiting hydrologic parameters were identified, results from the present study imply caution is needed in using the current WFD biological indicators based on analogous principles to the one adopted in Italy, especially to guide the management of ecological flows. Further research is needed to better quantify flow-ecology relationships and develop hydrology-sensitive indicators. Similar efforts were pursued by other countries where empirical waterflow preferences of benthic invertebrates were synthesised into a river flow index (e.g. LIFE index; Extence et al., 1999).

However, the validity of the LIFE index in other environmental settings needs to be tested (Dunbar et al., 2010) and, more generally, the index is designed to reflect changes in flow velocity and might correlate poorly with other flow parameters likely affected by river regulation (i.e. frequency and magnitude of variation). Ideally, effort and resources should be directed to the development of ecological indicators targeting specific flow characteristics that are most likely altered by river regulation and water uses.

## 6. Acknowledgements

This project has received funding from the European Union’s Horizon 2020 research and innovation programme under the Marie Skłodowska-Curie Grant Agreement No. 748969, awarded to SL. The authors want to thank the Environmental Agency of the Autonomous Province of Trento (APPA-TN) and the Environmental Agency of the Autonomous Province of Bolzano (APPA-BZ) for providing the STAR-ICMi, LIMECO and MQI datasets; the Hydro offices for providing the hydrological data; Elisa Stella (University of Trento) for providing the stream network and hydropower dams GIS data.

## SUPPLEMENTARY MATERIAL

### Appendix A

Description of the STAR_ICMi index (adapted from http://www.life-inhabit.it/cnr-irsa-activities/en/cnr-irsa-activities-related-inhabit/ecological-status/staricmi Buffagni et al., 2005, 2007, 2008).

The STAR_ICMi is a multimetric index which includes six metrics:

1. Average Score Per Taxon (Armitage et al., 1983);
2. Log10(sel_EPTD+1): Log10 (sum of Heptageniidae, Ephemeridae, Leptophlebiidae, Brachycentridae, Goeridae, Polycentropodidae, Limnephilidae, Odontoceridae, Dolichopodidae, Stratyomidae, Dixidae, Empididae, Athericidae & Nemouridae;
3. 1-GOLD 1:1 - (relative abundance of Gastropoda, Oligochaeta and Diptera);
4. Number of EPT families;
5. Total number of families
6. Shannon-Weiner diversity index.

The calculation of the STAR_ICMi is performed in 4 steps:

1. calculation of the raw value for each of the 6 metrics;
2. calculation of the Ecological Quality Ratio for each of the 6 metrics by dividing the observed value (i.e. obtained for the considered samples) by the median value of the metric calculated from the reference river type;
3. calculation of the weighted average of the EQR using a specifically assigned weight for each metric (0.333, 0.266, 0.067, 0.167, 0.083, 0.083, respectively for metrics 1-6);
4. normalization of the obtained value by dividing the value of the considered sample by the STAR_ICMi expected in reference samples.

Values of STAR_ICMi vary between 0 and +1, and different intervals correspond to the five quality classes defined by the Water Framework Directive: high, good, moderate, bad, poor.

## SUPPLEMENTARY FIGURES

**Supplementary Figure S1.**
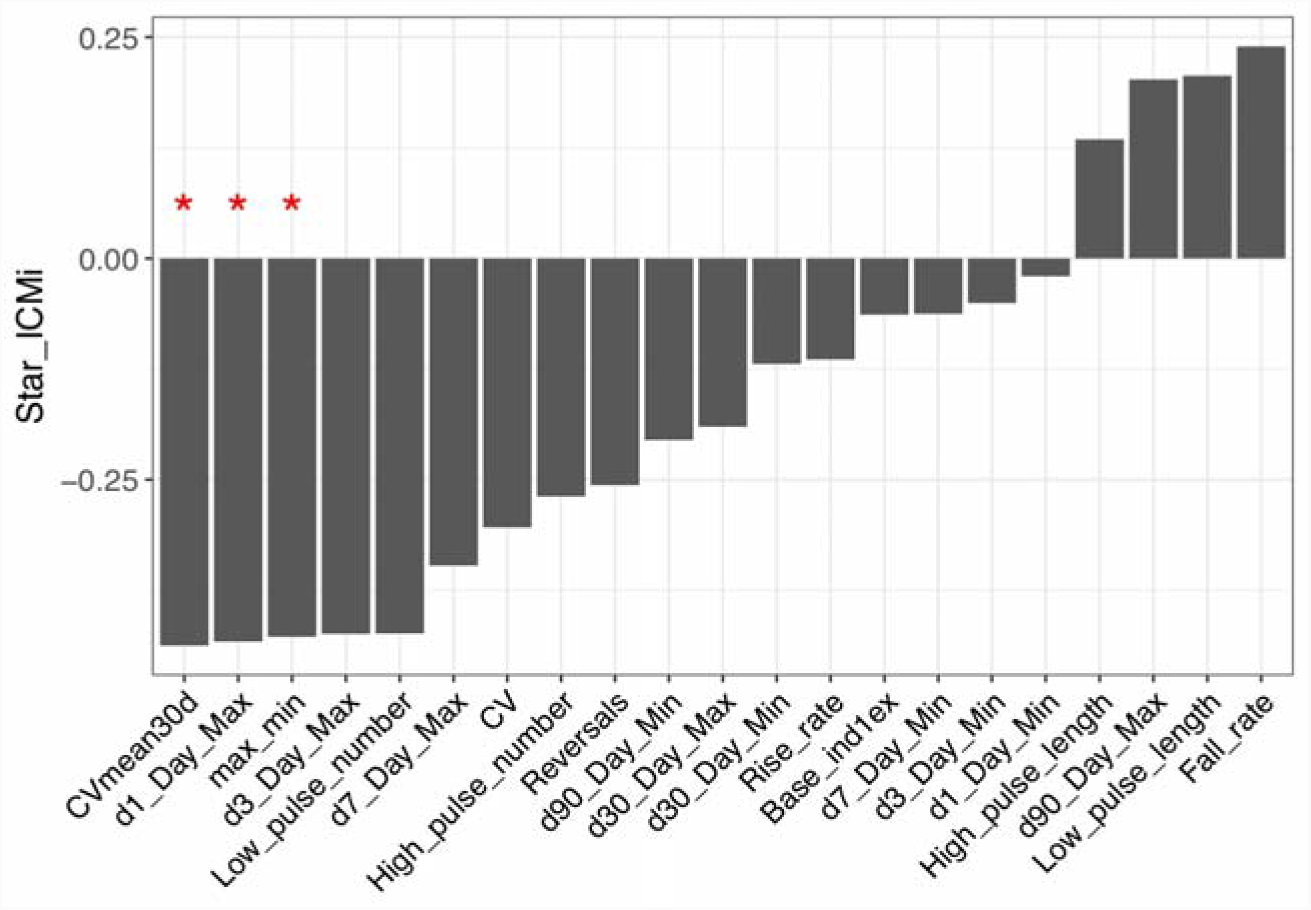
Correlation coefficients between the Star_ICMi and flow parameters calculated from 1-year flow-series preceding the biological sampling. Only three flow parameters showed significant correlation (red stars) based on linear mixed models.

**Supplementary Figure S2.**
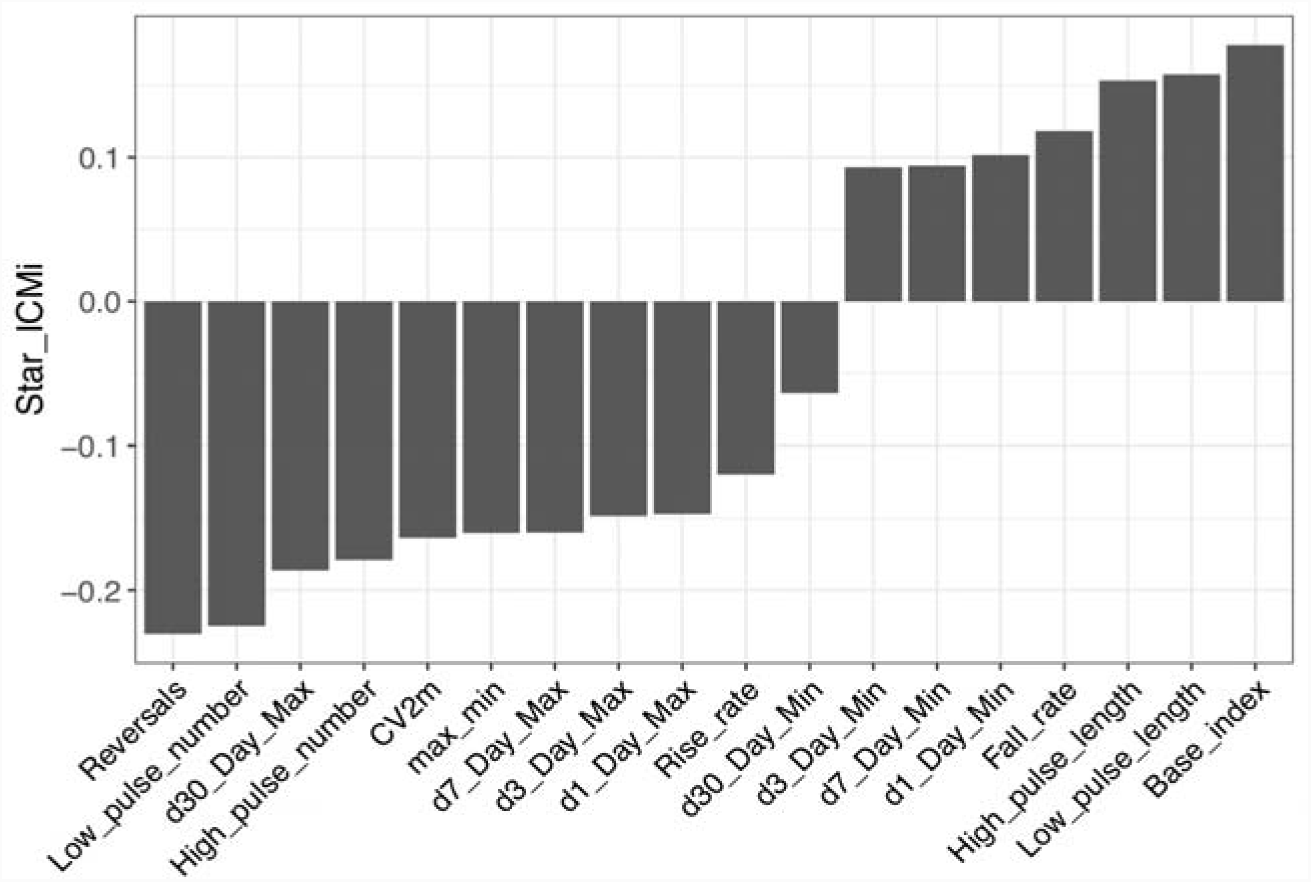
Correlation coefficients between the Star_ICMi and flow parameters calculated from 60-days flow-series preceding the biological sampling. No significant correlation was observed based on mixed models.

**Supplementary Figure S3.**
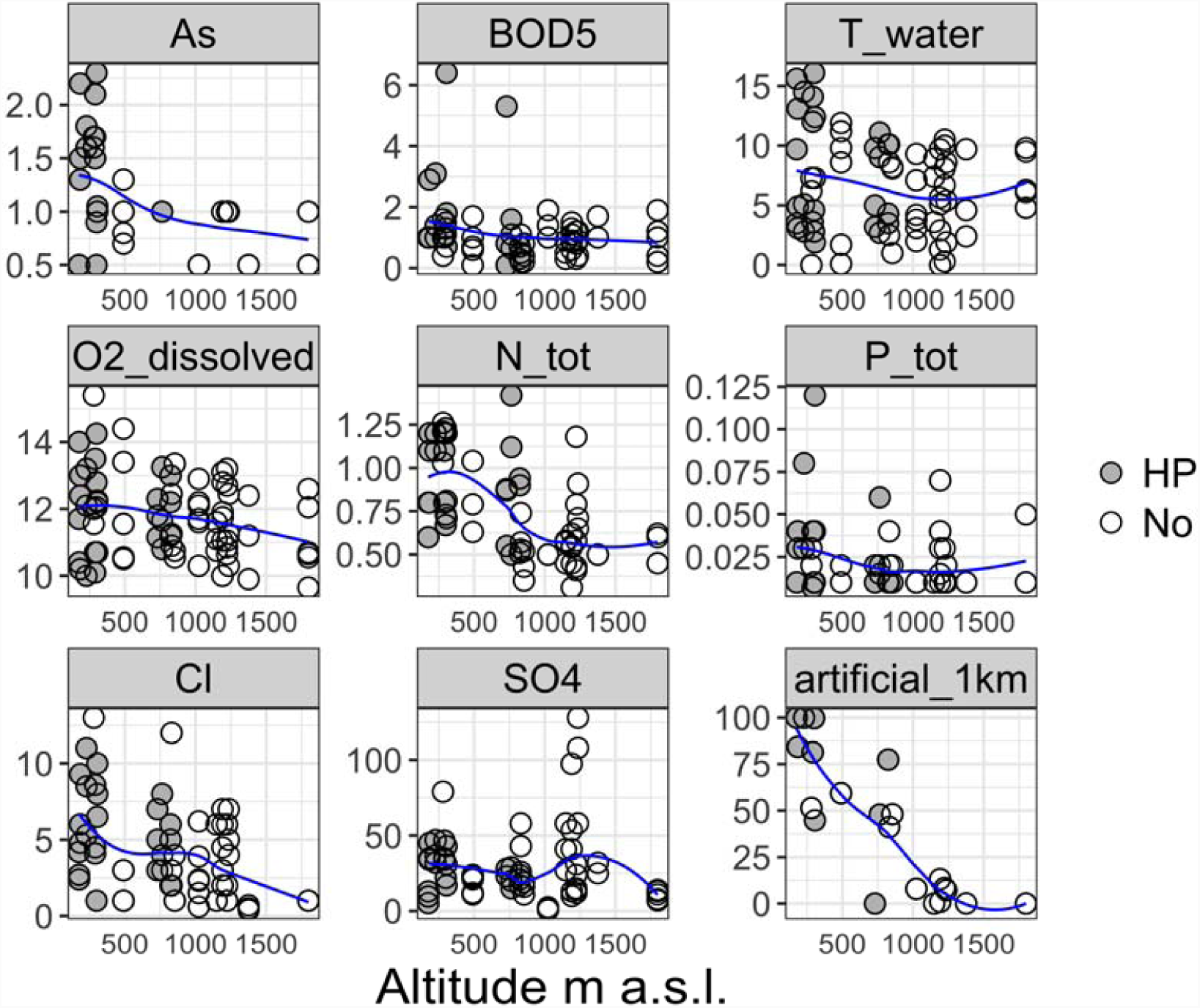
Relationship between altitude and main physico-chemical parameters and local land use across the study reaches used in the flow-ecology analysis. Grey circles = reaches affected by hydropower upstream (HP); white circles = reaches not affected by hydropower (No)

